# Aging and the Spectral Properties of Brain Hemodynamics

**DOI:** 10.1101/2024.12.05.626723

**Authors:** Ki Yun Park, Abraham Z. Snyder, Manu S. Goyal, Timothy O. Laumann, John J. Lee, Babatunde Adeyemo, Nicholas Metcalf, Andrei G. Vlassenko, Joshua S. Shimony, Eric C. Leuthardt

**Author notes:** **Corresponding Author:** Dr. Eric C. Leuthardt, Shi H. Huang Professor of Neurological Surgery, Washington University School of Medicine, Cortex 1 Building, 4320 Forest Park Ave, St. Louis, MO 63108.

## Abstract

Cerebral glucose metabolism (CMRGlc) systematically decreases with advancing age. We sought to identify correlates of decreased CMRGlc in the spectral properties of fMRI signals imaged in the task-free state. We analyzed lifespan resting-state fMRI data acquired in 455 healthy adults (ages 18-87 years) and cerebral metabolic data acquired in a separate cohort of 94 healthy adults (ages 25-45 years, 65-85 years). We characterized the spectral properties of the fMRI data in terms of the relative predominance of slow vs. fast activity using the spectral slope (SS) measure. We found that the relative proportion of fast activity increases with advancing age (SS flattening) across most cortical regions. The regional distribution of spectral slope was topographically correlated with CMRGlc in young adults. Notably, whereas most older adults maintained a youthful pattern of SS topography, a distinct subset of older adults significantly diverged from the youthful pattern. This subset of older adults also diverged from the youthful pattern of CMRGlc metabolism. This divergent pattern was associated with T2-weighted signal changes in frontal lobe white matter, an independent marker of small vessel disease. These findings suggest that BOLD signal spectral slope flattening may represent a biomarker of age-associated neurometabolic pathology.

**Significance Statement:** Aging is associated with a decline in cerebral glucose metabolism (CMRGlc). Here, we identified correlates of CMRGlc in the spectral properties of resting-state fMRI data using spectral slope (SS), which quantifies the relative predominance of slow vs. fast activity. We found that SS flattening with advancing age is most prominent in regions characterized by high CMRGlc in youth. A subset of older adults who diverged from the youthful pattern of SS topography also exhibited evidence of frontal lobe white matter pathology. These findings suggest that the spectral properties of rs-fMRI data provide mechanistic insights into age-related neuropathology. In particular, spectral slope measures may serve as early indicators of age-related neurometabolic decline and potentially identify individuals in whom intervention is indicated.

## Introduction

Brain metabolism and physiologic integrity decline with age in a manner parallel to numerous other measures (Kirkwood 2005, Lopez-Otin, Blasco et al. 2013). This principle has been demonstrated with Positron Emission Tomography (PET) (Kety 1956, Kuhl, Metter et al. 1982, Goyal, Hawrylycz et al. 2014) as well as resting-state functional magnetic resonance imaging (rs-fMRI) (Sala-Llonch, Bartres-Faz et al. 2015). In particular, cerebral blood flow and *oxidative* metabolism of glucose globally decline with advancing age (Goyal, Hawrylycz et al. 2014). In contrast, *glycolytic* metabolism of glucose declines most in regions with high values in youth (Goyal, Vlassenko et al. 2017, Goyal, Blazey et al. 2023). Thus, the regional pattern of cerebral metabolism in older adults often diverges from the normative youthful pattern (Goyal, Blazey et al. 2023). Intriguingly, some older adults retain youthful metabolic patterns while others do not (Goyal, Blazey et al. 2023).

Previous rs-fMRI studies of aging have largely focused on functional connectivity (FC) (Sala-Llonch, Bartres-Faz et al. 2015). FC refers to pairwise correlations of blood-oxygen-level-dependent (BOLD) signals in functionally related regions known as resting state networks (RSNs) (Biswal, Yetkin et al. 1995, Lowe, Mock et al. 1998, Cordes, Haughton et al. 2000, Hampson, Peterson et al. 2002, Greicius, Krasnow et al. 2003, Beckmann, DeLuca et al. 2005, Fox, Corbetta et al. 2006, Seeley, Menon et al. 2007). Comparatively fewer studies have evaluated age-related changes in the statistics of regional spontaneous BOLD signals. These studies have employed a variety of metrics including amplitude of low-frequency fluctuations (ALFF) and BOLD signal variability, both of which relate to the temporal variance of BOLD fMRI signals. Recent results have shown that developmental trajectories of ALFF in a cohort aged 8 to 18 years are topographically organized in relation to the sensorimotor-association cortex axis (Sydnor, Larsen et al. 2023). Additionally, aging studies have demonstrated reduced signal amplitude and less between-region variability in old as compared to young adults (Garrett, McIntosh et al. 2011, Grady and Garrett 2014). However, these previous reports have not considered the spectral characteristics of BOLD fMRI signals.

BOLD fluctuations are 1/f-like, i.e., characterized by an approximately linear decrease in log spectral power with increasing frequency (Bullmore, Long et al. 2001, He 2011, Tagliazucchi, von Wegner et al. 2013). Importantly, the spectral properties of resting-state BOLD signals vary across brain networks (Raut, Snyder et al. 2020), depend on the task state (He 2011), and change in patients with brain tumors and clinical comorbidities (<u>Park, Snyder et al.</u>). The spectral properties of BOLD signals also vary regionally in relation to brain glucose metabolism (Aiello, Salvatore et al. 2015, Nugent, Martinez et al. 2015, Deng, Franklin et al. 2022).

The preceding considerations suggest that the spectral characteristics of resting-state BOLD signal fluctuations may correspond with changes in cerebral blood flow and metabolism across the lifespan. To address this question, we evaluated the spectral properties of BOLD signal fluctuations and their correlations with cerebral blood flow and metabolism using two independent datasets: the Cambridge Centre for Ageing and Neuroscience (Cam-CAN) rs-fMRI dataset of 455 subjects aged 20-87 years (Shafto, Tyler et al. 2014, Taylor, Williams et al. 2017) and the Adult Metabolism & Brain Resilience (AMBR) metabolic dataset including 94 subjects aged 25-85 years (Goyal, Blazey et al. 2023).

## Results

We evaluated changes in the spectral slope (SS) of BOLD fluctuations across the cortex using a cross-sectional group of 455 individuals aged 20 to 87 years. SS was computed using linear regression of log power against frequency within the 0.015-0.145 Hz band. In general terms, this metric indexes the relative prevalence of slow vs. fast BOLD activity. Prior works have demonstrated a correlation between steeper spectral slopes, i.e., greater coherence at lower frequencies within the infraslow frequency range, and higher glucose metabolism (Aiello, Salvatore et al. 2015, Deng, Franklin et al. 2022). To characterize age-related SS changes for each cortical parcel, we fit parcel-specific, generalized additive models (GAMs), parametric in age, with sex and head motion as linear covariates, as previously described in (Sydnor, Larsen et al. 2023). Each GAM estimates the parcel-wise trajectory of changes in SS with age.

SS flattens with increasing age. In **Figure 1**, panel A shows BOLD fluctuations sampled from the same ROIs in two subjects in both time and frequency domains. Panel B illustrates the average power spectra, computed by averaging across all cortical voxels and participants within each age group. Panel C shows the topography of SS change from youth to old age. Warmer hues indicate a steeper SS, reflecting a greater prevalence of slow over fast activity within the infra-slow frequency range. The steepest spectral slopes were observed in medial and lateral parietal, prefrontal, and visual cortices. This topography remained consistent across all age groups, suggesting that, despite a general flattening observed in the brain, major features of SS topography are generally preserved in most older individuals.

**Figure 1.**
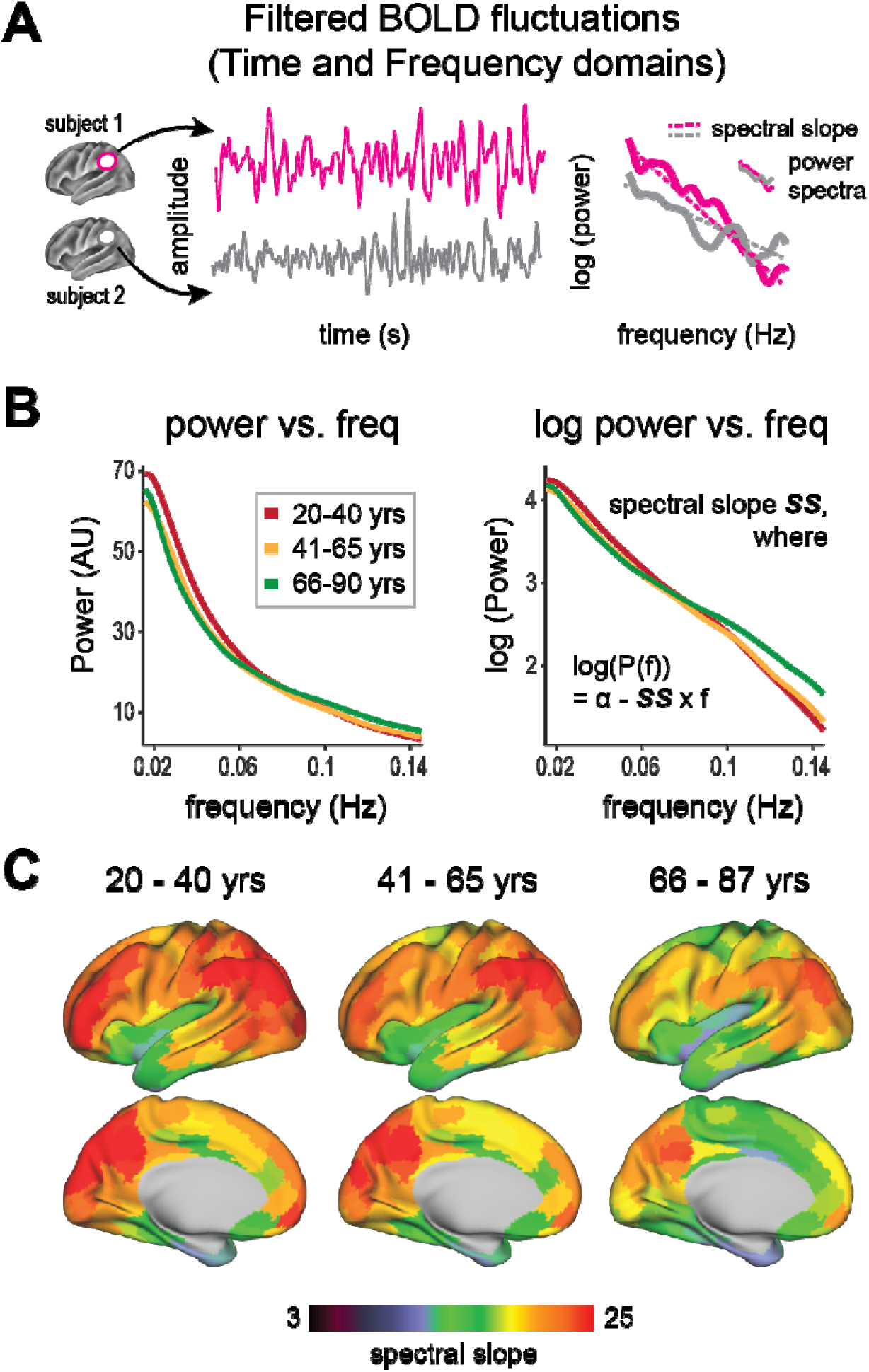
Spectral slope (SS) flattens with advancing age. **A.** Two distinct BOLD fluctuations, sampled from the same ROIs, in two exemplar subjects, shown in both time and frequency domains. **B. (left)** Averaged power spectra across participants in three age groups. **(right)** Averaged power spectra plotted as log power vs. frequency. SS is defined as the negative of the first derivative of the slope fitted to the log power. **C.** Parcellated surface SS maps (footnote: the volumetric data was projected onto the Conte-69 average inflated surface for surface-based visualization), averaged across participants within three age groups, illustrate the topography of age-related SS flattening across the lifespan. Cortical regions with steep spectral slopes include medial and lateral parietal, prefrontal, posterior cingulate, and visual cortices (warmer hues). The age cohorts are 20-40 years (n = 113), 41-65 yrs (n = 179), and 66-87 yrs (n = 163). Although age-related SS flattening differs across cortical regions, the characteristic features of SS topography persist across all age groups.

Visualization of regional fits with significant age-related effects suggested that there were two distinct trajectories of spectral slope behavior across the lifespan: 1) consistent flattening of spectral slope throughout life, or 2) no decline until mid-adulthood, followed by marked and continuous flattening. Fuzzy c-means clustering confirmed the presence of these varying trajectories across cortical parcels. Silhouette score analysis (Rousseeuw 1987) indicated that two clusters were optimal (see **Supplemental Materials**). Whereas most cortical regions exhibited continuous SS flattening (**Fig. 2A**), certain regions, particularly those coinciding with auditory (AUD), cingulo-opercular (CON), and salience networks (SAL), exhibited flattening starting in mid-adulthood (**Fig. 2B**). Regions within Cluster 2 include dorsal anterior cingulate cortex (dACC), frontal operculum, supplementary motor area (SMA), supramarginal gyrus (SMG), inferior frontal gyrus (IFG), pars marginalis of the cingulate gyrus, and the mesial surface of the visual cortex.

**Figure 2.**
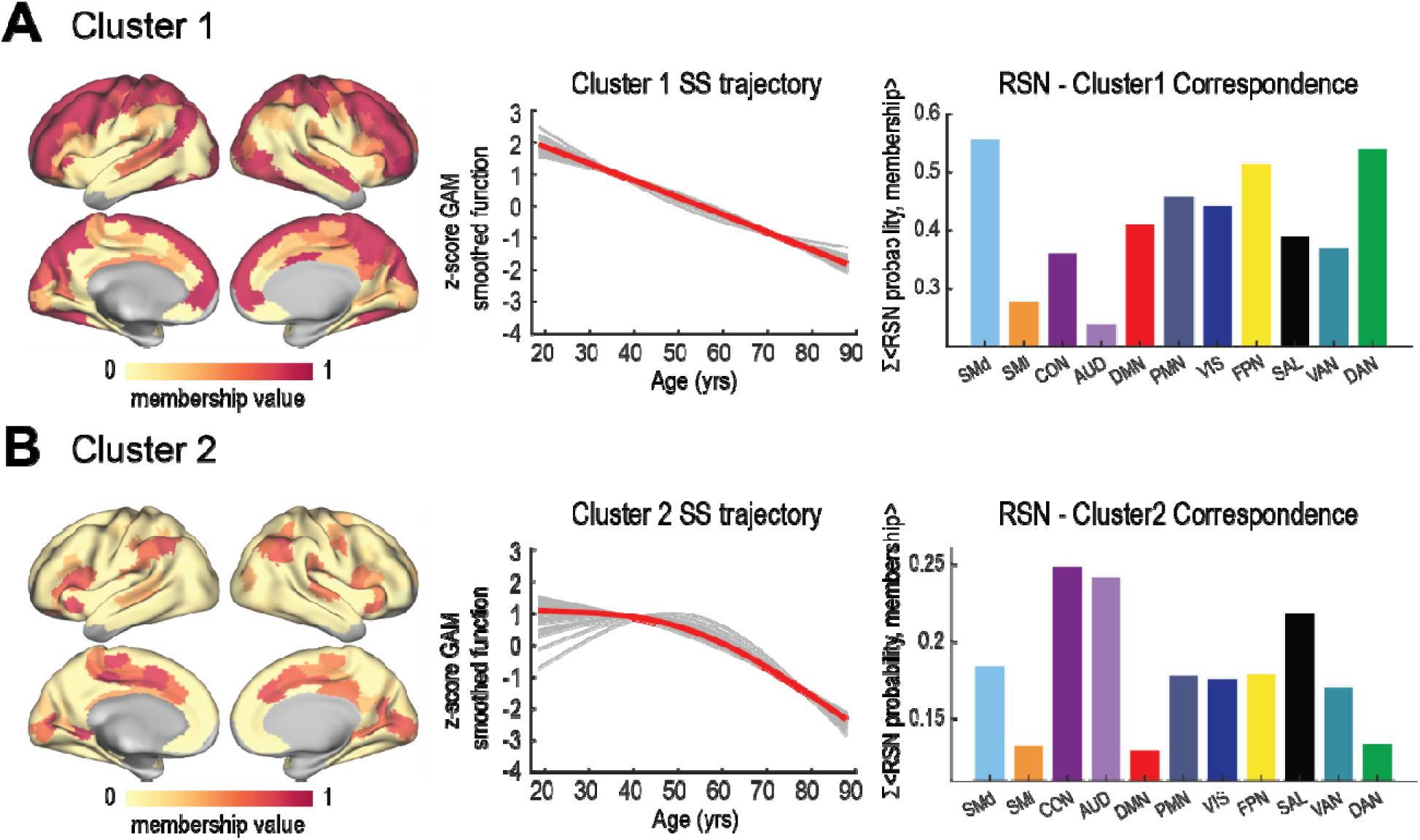
Trajectories of age-related SS flattening across cortical parcels (Schaefer 300 parcellation scheme), identified by fuzzy c-means clustering. **A.** Cluster 1, comprising the majority of cortical parcels, exhibits a consistent flattening of the spectral slope with age. The right-most panel displays the “RSN-Cluster1 Correspondence Index,” computed as the normalized sum of dot products between RSN probability maps (Luckett, Lee et al. 2022) and membership values for Cluster 1. **B.** Cluster 2, comprising cortical parcels primarily within the auditory, cingulo-opercular, and salience networks, exhibits delayed flattening that begins in mid-adulthood (∼50 yrs). The “RSN-Cluster2 Correspondence Index” is shown on the rightmost panel. Clustering analysis was done using the Schaefer 300 parcellation scheme in a volumetric space; results are subsequently projected onto a surface for visualization. Parcels heavily affected by susceptibility related signal loss (Ojemann, Akbudak et al. 1997)) or insignificant age-related effects were excluded in this analysis.

Significant SS changes were observed in 82% of cortical parcels (p_FDR_ < 0.05). The regional magnitude of age effects was assessed using age-specific R^2^ difference, which measures the differences in explained variance between the full model (with age) and the reduced model (without age) (**Fig. 3A**). Age-specific R^2^ difference varied from 0 to 0.12, where 0 indicates that SS change is attributable to covariates other than age. High age-specific R^2^ difference values denote regions where age heavily contributed to SS change. Significant age effects were found in lateral and medial prefrontal regions, superior parietal lobule, intraparietal sulcus, the posterior cingulate cortex, and the medial surface of the visual cortex. Interestingly, age-specific R^2^ difference topography markedly resembled the distribution of CMRGlc across the cortical regions (**Fig. 3B, C**; Spearman correlation ρ = 0.66, spin test p value (p_spin_) = 3e-04). Brain regions with higher levels of glucose metabolism in youth showed greater effects of age on SS (**Fig. 3C**).

**Figure 3.**
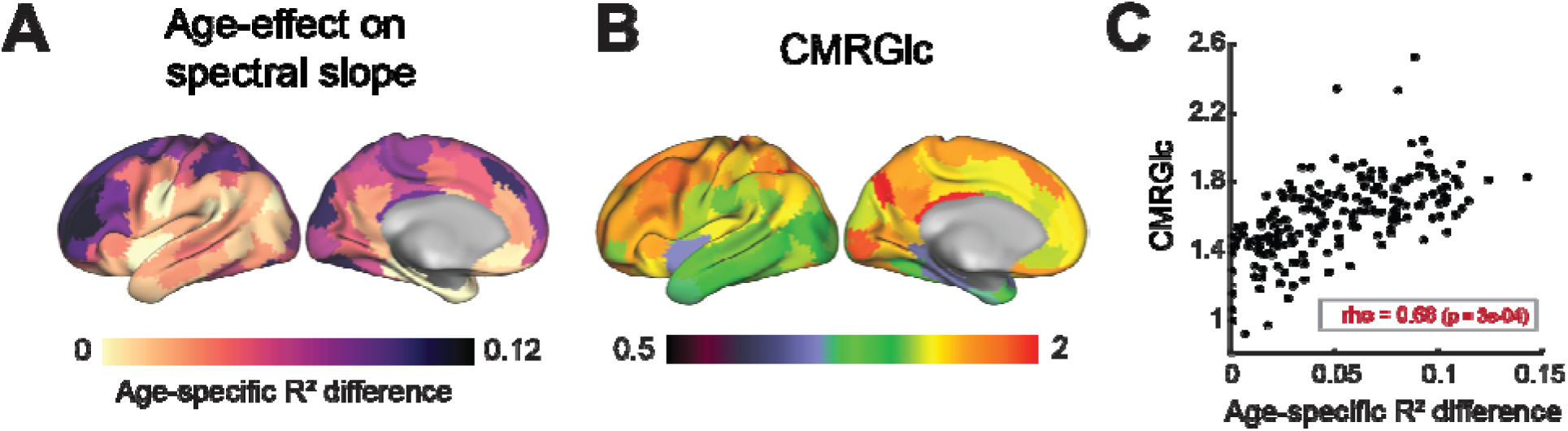
**A.** Effect of age on SS (age-specific R^2^ difference) across the cortical surface. **B.** Parcellated cerebral metabolic rate of glucose use (CMRGlc) map averaged across 30 healthy young adults aged 25-45 years (AMBR dataset from (Goyal, Blazey et al. 2023)). **C.** Spearman correlation between age-specific R^2^ difference and the average CMRGlc map (rho = 0.66, p = 3e-04). Each symbol represents one parcel. Both metrics were evaluated using Schaefer 200 parcellation scheme in a volumetric space and subsequently projected onto the surface for visualization.

With increasing age, cerebral glucose metabolism not only decreases quantitatively (Kuhl, Metter et al. 1982, Dastur 1985, Goyal, Vlassenko et al. 2017) but also loses its youthful regional patterns in some – but not all – older adults (Goyal, Blazey et al. 2023). Considering the established association between the spectral properties of BOLD signals and CMRGlc (Aiello, Salvatore et al. 2015, Nugent, Martinez et al. 2015, Deng, Franklin et al. 2022), and based on our present finding that SS flattens with aging, we asked whether SS topography – its inherent youthful regional variations in spectral slope – also loses its youthful pattern with increasing age. We assessed this by evaluating the SS Youthful Index (SSYI), evaluated as the Spearman correlation between individual SS maps and the averaged SS_young_ map (see **Methods 7.1**; **Fig. 4A**).

**Figure 4.**
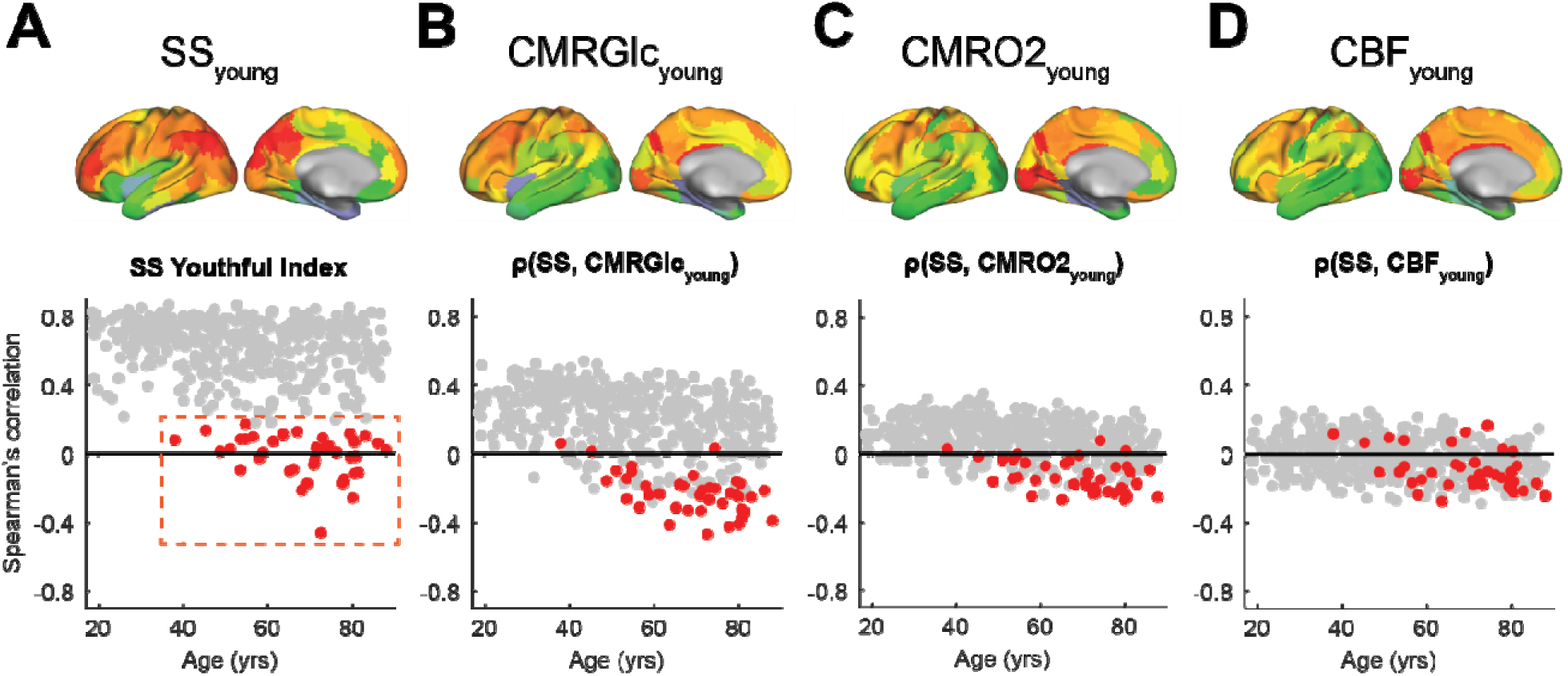
**(Top)** Topography of average spectral slope (SS) maps, CMRGlc, CMRO_2_, and CBF of the younger cohort (25-45 yrs). SS_young_ map was computed by averaging SS maps of younger participants (<45 yrs) in the CamCAN dataset. CMRGlc, CBF, and CMRO_2_ maps are from the AMBR dataset (see **Methods 1.2.**). **(Bottom)** Each symbol (grey or red) represents an individual’s Spearman correlation between their SS map and CamCAN-average SS_young_ map (panel **A**, defined as SS Youthful Index). Spearman correlations between SS maps and AMBR-average CMRGlc_young_, CMRO2_young_, and CBF_young_ maps are shown in panels **B-D**. Red symbols identify outlier individuals whose correlation values are greater than 3 scaled median absolute deviations from the median. These individuals also show negative and weaker correlations with CMRGlc_young_ and CMRO2_young_ maps. Correlations between SS maps and CBF_young_ and CMRO2_young_ are weaker as compared to those with CMRGlc_young_. SS Youthful Indices (panel **A**) and correlations between CMRGlc_young_ and SS maps (panel **B**) are generally positive in younger CamCAN participants. However, an increasing fraction of individuals diverge from the youthful pattern starting at age ∼45.

Notably, the youthful pattern of SS topography generally persisted across the lifespan. However, SSYI inter-subject variability increased cross-sectionally with advancing age. Importantly, a distinct subset of older individuals displayed significantly weaker or even negative values of SSYI (less than three scaled median absolute deviations from the median). We labeled these individuals as outliers (red symbols in **Fig. 4A**) for further analysis. Interestingly, these outliers also exhibited weaker correlations with CMRGlc_young_ and CMRO2_young_ (**Figs. 4B-C**). This trend appeared much less distinguishable when correlated with CBF (**Fig. 4D**).

Considering that older subjects typically exhibit greater grey matter atrophy and increased head motion, we hypothesized that the identified outliers (red symbols in **Fig. 4A**) would demonstrate smaller cortical volume and/or more head motion. Moreover, as previous findings demonstrated that the adult female brain is metabolically more “youthful” (<u>Goyal, Blazey et al.</u>), we assessed the effect of sex on the SS Youthful index in outliers and non-outliers aged over 45 years. No significant differences were observed in grey matter volume, head motion, or sex between these groups. In **Fig. 5A-B**, the overlays of black symbols on red symbols indicate comparable distributions of grey matter volume and head motion in both groups. The histogram in **Fig. 5C** shows no clear dominance of either sex in either group.

**Figure 5.**
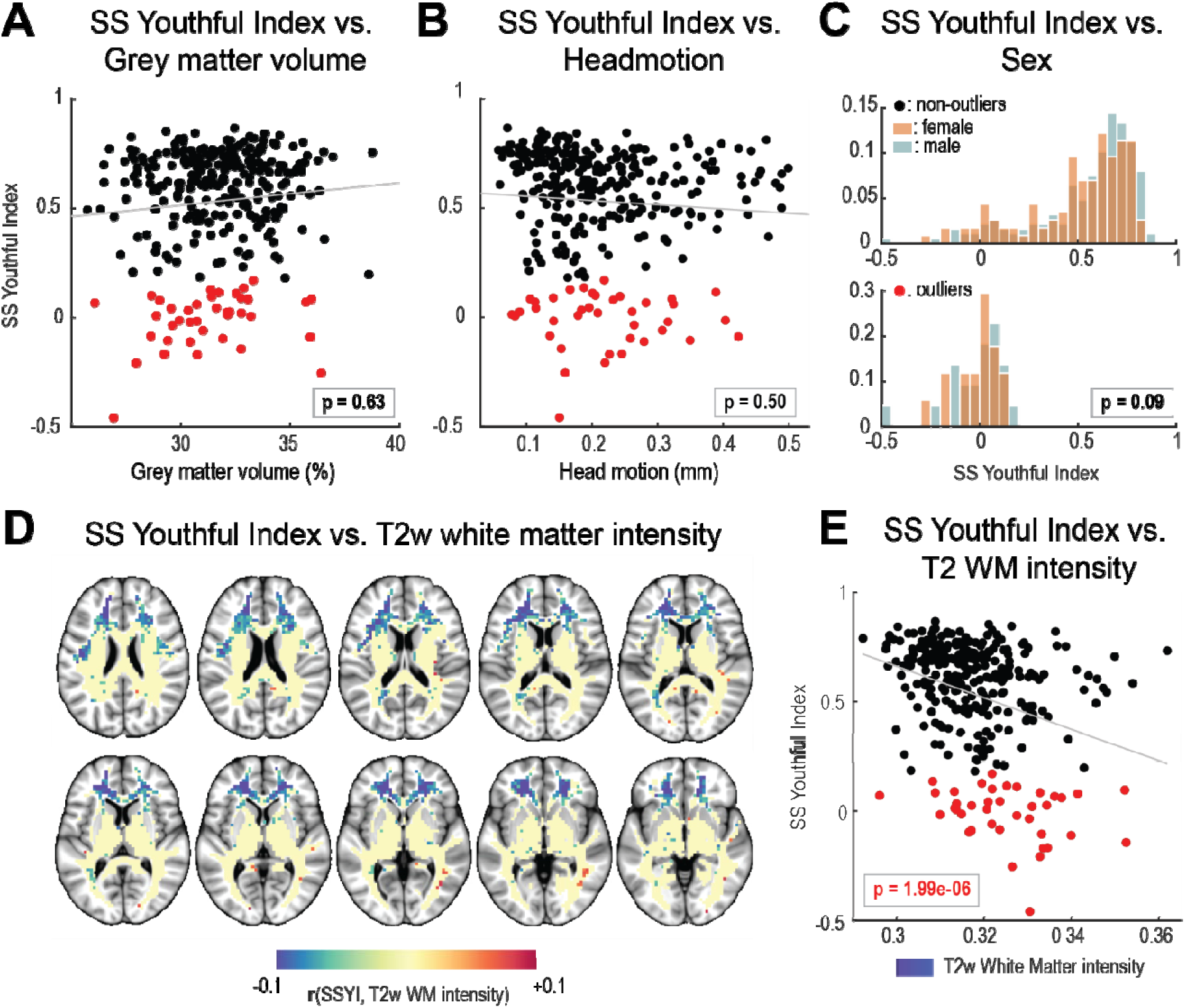
SS Youthful Index is significantly negatively associated with frontal lobe white matter (FLWM) T2w image intensities. SS Youthful Index is not associated with grey matter volume, head motion, and sex. A group of older subjects (age > 45yrs) was selected to determine if the outliers identified in Fig. 4 exhibited significant differences in their grey matter volume, head motion, sex, or WM T2w image intensities (see **Methods 7.3**). Outliers (red symbols in Fig. 4) are shown as red symbols. Non-outliers (grey symbols in Fig. 4) are shown in black. **A.** Grey matter volume vs. SS Youthful Index. **B.** Head motion (mm) vs. SS Youthful Index. **C.** Differences in sex distributions between the two groups: non-outliers (top, black symbols in Fig. 5A**-B**) and outliers (bottom, red symbols in **Figs. 5A-B**). **D.** Voxelwise correlations across participants between normalized T2w signal intensity and SSYI (as in voxel-based lesion-symptom mapping (VLSM) (Bates, Wilson et al. 2003)). Significant Pearson correlations (p <0.05) are shown in blue. **E.** Averaged T2w signal intensity within the blue-shaded regions in panel D vs. SSYI. Statistical significance was evaluated in terms of linear regression model fit of age, gray matter volume, head motion, sex, and frontal lobe WM T2w image intensity as predictors of SS Youthful Index outlier status. The p-values shown in panels A-C and E are from the fit linear regression model result (see Supplementary Table 1).

Intriguingly, however, SS Youthful Index was associated with higher T2-weighted image intensities, predominantly in the frontal lobe white matter (FLWM; left panel in **Fig. 5D**). This focal topography was identified by correlating participants normalized T2w image intensities with SS Youthful Indices in each voxel (see **Methods 7.3**). The right panel of **Fig. 5D** illustrates the relation between SSYI and FLWM T2w intensity values in the blue-shaded regions of FLWM.

To quantify these observations, we fit a linear regression model using age, grey matter volume, head motion, FLWM T2w image intensities, and sex as predictors, and the binarized SS Youthful Index as the response variable. The only statistically significant predictor was FLWM T2w image intensity (estimate = −7.40, p-value = 1.99e-06). This result suggests that, among the middle-aged to elderly subjects, deviations in the SS Youthful Index are more associated with frontal lobe white matter abnormalities than grey matter atrophy, head motion, sex, or age (**Supplementary Table 1**). FLWM T2w image intensities were also significantly positively correlated with body mass index (BMI) (**Supplementary Figure 3**).

## Discussion

As the global population ages, there is a pressing need for accessible whole-brain imaging biomarkers of functional integrity of the brain. Here, we propose that the spectral properties of rs-fMRI BOLD signals may provide a functional biomarker of pathologic brain aging. fMRI has historically been primarily used for mapping task-based regional changes or identifying functional topographies using resting state functional connectivity (RSFC) (Van Essen, Ugurbil et al. 2012). The current work focuses on a complementary approach that evaluates age-related changes in BOLD signal spectra, with a particular focus on their relations with cerebral metabolism.

Cerebral metabolism declines with age (Goyal, Vlassenko et al. 2017). We found that resting-state BOLD fMRI signal fluctuations typically exhibit progressive spectral slope (SS) flattening with age, most prominently in cortical regions with high levels of glycolytic metabolism in younger individuals. These same regions also exhibit age-associated loss of aerobic glycolysis (Goyal, Blazey et al. 2023). Thus, spectral flattening appears to be associated with loss of aerobic glycolysis. Consistent with prior work, we demonstrate a topographical correlation between the spectral characteristics of BOLD fluctuations and CMRGlc (Aiello, Salvatore et al. 2015, Nugent, Martinez et al. 2015, Deng, Franklin et al. 2022), which is most apparent in younger adults. Additionally, the present results suggest that while most older individuals maintain a youthful SS topography (as evaluated using SSYI), a subset of older individuals deviates significantly from the youthful pattern. Notably, lower SSYI is associated with higher T2-weighted image intensities in frontal lobe white matter. Increased T2-weighted signal intensity is a marker of white matter disease (Liu, Yang et al. 2017), previously shown to be particularly prominent in older individuals in the same regions highlighted in **Fig. 5D** (Salat, Kaye et al. 1999, Bartzokis, Cummings et al. 2003). These observations suggest that SSYI correlates with other biomarkers of pathological neurometabolic aging.

### Analyzing BOLD fluctuations in the temporal frequency domain

The physiological significance of spectral slope flattening follows from the notion that infraslow frequencies (nominally, < 0.1Hz) primarily reflect signals of neural origin (Palva and Palva 2012, Mitra and Raichle 2016), whereas faster frequencies represent artifact, primarily thermal noise, and head motion (Liu 2016). As head motion manifests as intermittent burst noise, its spectrum is theoretically white, as is that of thermal noise. In contrast, direct measurements show that BOLD fMRI signals are very attenuated at frequencies greater than 0.2 Hz (Boynton, Engel et al. 1996). Thus, BOLD power spectra generally assume the characteristic of a 1/f-like spectrum superimposed on a flat noise floor. Hence, the spectral slope (SS) indexes the extent to which BOLD fMRI fluctuations are of neural-as opposed to non-neural origin. Empirically, in glioblastoma patients, an increased proportion of high vs. low-frequency rs-fMRI BOLD activity is associated with compromised brain integrity (Park, Snyder et al. 2023). Here, we extend this principle to aging.

### Spectral slope flattening in aging

The underlying physiology defining the spectral slope of the BOLD signal, as well as its changes, are as yet uncertain. Despite this limited understanding, prior work has shown that the spectral properties of rs-fMRI signals depend on task state (He 2011), correlate with cerebral glucose metabolism (Aiello, Salvatore et al. 2015, Nugent, Martinez et al. 2015, Deng, Franklin et al. 2022), and are functionally organized (Raut, Snyder et al. 2020). Critically, regions with steeper spectral slopes exhibit stronger functional connectivity across extensive regions (Raut, Snyder et al. 2020) and spatially correspond with higher baseline CMRGlc (**Fig. 4B**). This circumstantial evidence suggests a potential relation between the required metabolic supply and the maintenance of cellular or synaptic dynamics (Goldman, Compte et al. 2009) engaging in large-scale circuits over multiple timescales. Aging affects this process, perhaps owing to the simplification of neuronal architecture (e.g., reduced arborizations and dendritic length, decreased spine numbers (<u>Dickstein, Kabaso et al.</u>), accumulation of DNA damage and mutations (<u>Lombard, Chua et al.</u>), or inefficiencies in DNA repair processes (<u>Gorbunova, Seluanov et al.</u>). Accordingly, it is plausible that age-related neuropathology manifests as spectral slope flattening of BOLD signals.

Importantly, age-associated spectral slope flattening is more pronounced in regions with higher CMRGlc in young adults (**Fig. 3**). The pathophysiological mechanisms underlying this relation are incompletely understood. It is known that Aβ deposition is a correlate of neural activity in the APP mouse model of Alzheimer disease (Bero, Yan et al. 2011). Moreover, Aβ deposition is more prevalent in parts of brain that are more strongly functionally connected with other parts of the brain (i.e., increased global FC magnitude), both in the APP mouse (Bero, Bauer et al. 2012) and healthy humans (<u>Buckner, Sepulcre et al.</u>). These observations suggest a potential “burn-out” mechanism for neural integrity over the lifespan. The relevant stressor may not be single unit discharge, as action potentials are energetically inexpensive (Raichle and Mintun 2006).

Speculatively, the relevant stressor may be limits on the extent to which synaptic reweighting is able to preserve previously learned memories while simultaneously encoding new information (Tononi and Cirelli 2014, Laumann and Snyder 2021). This would be consistent with the observation that sustained neural plasticity over a lifetime is associated with loss of synaptic spines and dendritic simplification (Dickstein, Kabaso et al. 2007). The preceding considerations suggest that these histological findings should be especially prevalent in individuals here identified as SS outliers. Testing this hypothesis would require a study combining ante-mortem rs-fMRI with post-mortem histology.

### Delayed onset of SS flattening exhibits regional specificity

While the majority of cortical regions showed continuous flattening of SS with increasing age, a subset of regions exhibited a delayed onset of SS flattening starting in mid-adulthood (**Fig. 2**). Despite their unique and varied functions, many of these regions share a common role in initiating and controlling goal-directed behaviors. Specifically, dACC, frontal operculum, and SMG are key components of the cingulo-opercular network (CON), associated with initiating, maintaining, and controlling task sets (Dosenbach, Fair et al. 2007, Dosenbach, Fair et al. 2008, Dosenbach, Raichle et al. 2024). SMA and IFG (includes Broca’s area) are involved in initiating and controlling movements and expressive language (Nachev, Kennard et al. 2008, Liakakis, Nickel et al. 2011). The dorsal PCC is functionally distinct from the DMN and has been proposed to be involved in both detecting and responding to changes in the environment (Leech and Sharp 2014). The delayed onset of changes in these brain regions may indicate a decline in the control of complex motor behaviors in older age, as individuals become more sedentary.

### SS Youthful Index progressively separates outliers from the “normal” aging group with increasing age; this separation correlates with frontal lobe white matter changes

The youthful pattern of spectral slope topography is generally maintained across the lifespan in most individuals. However, a subset of older individuals showed weaker and even negative SS Youthful Index values (**Fig. 4A**). Although all participants included in this study are cognitively unimpaired (as defined by MMSE score greater than 24) (<u>Taylor, Williams et al.</u>), cognitive normality does not exclude the possibility of asymptomatic neuropathology. In fact, the appearance of abnormal biomarker values often precedes the onset of clinical symptoms by several years (<u>Fagan, Xiong et al., Chan, Krebs et al.</u>). Furthermore, studies of normal aging have been shown to be confounded by failure to exclude participants positive for preclinical biomarkers (e.g., CSF biomarkers including Aβ42 and tau) (<u>Brier, Thomas et al.</u>). Hence, it is plausible that the present outlier subset in our sample harbors covert neuropathology.

Notably, SS Youthful Index was independent from grey matter volume or head motion, and differences in sex (**Figs. 5A-C**). However, we identified a significant negative correlation between SS Youthful Index and frontal lobe white matter (FLWM) T2w image intensities (**Fig. 5D**). This finding echoes previous work that linked white matter hyperintensity burden to the loss of the youthful pattern of brain aerobic glycolysis (<u>Goyal, Blazey et al.</u>). Additionally, aged white matter is more vulnerable to neurological disorders (Liu, Yang et al. 2017), and significant changes in FLWM have been observed as correlates of aging and AD (Salat, Kaye et al. 1999, Bartzokis, Cummings et al. 2003). Hence, it is conceivable that macrostructural frontal lobe white matter damage could contribute to a greater decline in neural function related to both aging and metabolic health.

Moreover, we found a significant positive correlation between FLWM T2w image intensities and body mass index (BMI) (see Supplementary Materials; **Fig. S4**). Whole brain spectral slope (averaged over parcels) and SSYI also were also independently associated with BMI after accounting for age, sex, and head motion as covariates (**Fig. S3**, **Tables S2** and **S3**). White matter hyperintensities are known to be associated with hypertension (Dufouil, de Kersaint-Gilly et al. 2001, Wardlaw, Smith et al. 2013, Rashid, Li et al. 2023) and BMI, a crude measure of body fat percentage, which is often regarded as a risk factor for cardiovascular disease (Brown, Higgins et al. 2000). Together, we hypothesize that systemic metabolic health may impact brain structure and function.

### rs-fMRI spectral slope: a potential biomarker of neuropathology

We propose a conceptual framework wherein biomarkers of neuropathology are characterized by change points at which the range of observed values progressively increases with age. In some cases, pathology leads to decreased values (e.g., hippocampal volume (<u>Luo, Agboola et al.</u>) or cortical thickness (<u>Pereira, Svenningsson et al., Walhovd, Storsve et al.</u>). In other cases, pathology leads to increased values (e.g., white matter hyperintensities (Luo, Ma et al. 2023), Aβ deposition (<u>Luo, Agboola et al.</u>), and accumulation of p-tau (<u>Luo, Agboola et al.</u>). In either case, the variance of biomarker values increases with age: some individuals maintain a normative or youthful state, whereas others deviate significantly from the youthful pattern. Within this framework, we hypothesize that the spectral slope of rs-fMRI BOLD signals is a potential accessible biomarker of pathological neurometabolic aging. The underlying mechanisms that lead to changes in the spectral characteristics of BOLD signal fluctuations remain to be determined.

### Limitations

We discuss several limitations of this work. BOLD signals are inherently sensitive to vascular and respiratory factors, as well as head motion, which is more prevalent in aging populations. Despite stringent quality control measures and accounting for head motion as a covariate, these factors may still affect the current findings. Additionally, partial volume correction (PVC) was applied in the AMBR dataset but not in the CamCAN dataset. PVC is particularly important in PET studies owing to the poor spatial resolution of older PET images (5-6 mm full-width half maximum) and is likely to have a lesser impact on fMRI images. However, it is still conceivable that the increased inter-subject variability observed in correlations between SS topographies and youthful metabolic maps may be attributed, at least in part, to a greater partial volume effect with aging. To evaluate white matter abnormalities, we used T2-weighted images, implementing a technique previously described by (Brier, Snyder et al. 2021), which normalizes each participant’s T2w image intensities to a reference intensity atlas. Future studies would benefit from using FLAIR to more accurately quantify abnormal white matter hyperintensities. Finally, the cross-sectional nature of our study limits our ability to track within-individual changes in BOLD fluctuations with aging. Longitudinal studies could provide more direct evidence of co-occurring age-related changes in metabolism and BOLD spectral slope.

## Materials and Methods

### 1. Datasets

#### 1.1. Cam-CAN dataset

The study included 455 participants from the adult lifespan Cambridge Centre for Ageing & Neuroscience (CamCAN) dataset (Shafto, Tyler et al. 2014, Taylor, Williams et al. 2017). Exclusion criteria included suboptimal registration due to compromised structural scans or significant head motion (averaged changes in head displacement and rotation in quadrature > 0.5mm). The demographics of our sample include an age range of 18 to 88 years (54.7 ± 18.5 yrs), comprising 276 males and 179 females. The recruitment and selection processes of the study participants are detailed in (Shafto, Tyler et al. 2014). Briefly, inclusion criteria included cognitively healthy subjects without communication or mobility issues, substance abuse problems, and those eligible for MRI scans. MRI datasets were collected at a single site (MRC-CBSU) using a 3T Siemens TIM Trio scanner with a 32-channel head coil. T1w and T2w structural images and resting state scans were used in our analysis. The resting-state fMRI data included an 8-minute 40-second run (261 frames) acquired while participants rested with their eyes closed. A T2*-weighted echoplanar imaging (EPI) sequence was used to collect 261 volumes, each containing 32 axial slices with slice thickness of 3.7mm, interslice gap of 0.74 mm for whole brain coverage (TR = 1970ms; TE = 30ms; flip angle = 78°; field of view (FOV) = 192×192mm; voxel-size = 3×3×4.44mm) (Taylor, Williams et al. 2017).

#### 1.2. Aging Metabolism & Brain Resilience (AMBR) dataset

Cognitively unimpaired adults without evidence of brain Alzheimer’s Disease (AD) pathology were recruited from Washington University in St. Louis community and the Knight Alzheimer Disease Research Center. A detailed explanation of the full dataset is in (Goyal, Blazey et al. 2023). Data from a total of 94 participants from this dataset were selected and divided into two groups: 30 young adults (ages 25-45 years; 16 males, 14 females); 64 older adults (ages 65-85yrs; 32 males, 32 females). All participants had undergone Positron Emission Tomography (PET) imaging using a Siemens ECAT HR+ scanner. The protocol included a single 18F-FDG scan, following a slow intravenous injection of ∼5 mCi FDG, and two sets of ^15^O PET scans. The final 20 minutes (40-60 minutes post-injection) of dynamic acquisition of the FDG images were summed and converted to SUVR measurements relative to the whole brain to assess the cerebral metabolic rate of glucose (CMRGlc). Each ^15^O PET session consisted of two sessions of three scans to measure cerebral blood volume (CBV), cerebral blood flow (CBF), and cerebral metabolic rate of O_2_ (CMRO_2_). Participants were instructed to remain awake with their eyes closed throughout the scans. Please see (Goyal, Blazey et al. 2023) for full details regarding data acquisition and preprocessing methods.

### 2. Data Acquisition and Preprocessing

#### 2.1. Cam-CAN dataset

Initial fMRI preprocessing followed the conventional practice (Shulman, Pope et al. 2010). Briefly, this included compensation for slice-dependent time shifts and rigid body correction of head movement within and across runs (Power, Barnes et al. 2012). The preprocessed data were then resampled to align the structural data in (3mm)^3^ atlas space using a composition of initial affine transform and a warping map (computed using the Advanced Normalization Tools (ANTs) registration) connecting the fMRI volumes with the T1w structural image. Details of the steps regarding ANTs registration are outlined in (Park, Snyder et al. 2023). Motion correction was included in the final resampling to generate volumetric timeseries in (3mm)^3^ atlas space. Further preprocessing steps included the removal of voxel-wise linear trends from each fMRI run, temporal low-pass filtering to retain frequencies below 0.15Hz, and regression of nuisance waveforms. These regressors included six head motion timeseries, average signals from CSF regions, and the signal evaluated over the whole brain (global signal regression). Finally, spatial smoothing was applied (6mm full width at half maximum Gaussian blur in each direction).

#### 2.2. Adult Metabolism & Brain Resilience (AMBR) dataset

To accurately estimate metabolic activity specific to a tissue type, partial volume correction (PVC) is essential owing to the limited spatial resolution of PET scans. The necessity arises from both “tissue fraction” and “point spread function” effects (Sattarivand, Kusano et al. 2012). Here, we employed a parcel-based approach for PVC, using Schaefer’s 200 parcellation scheme (detailed in the Methods Parcellations section), presuming that each parcel exhibits a distinct and uniform activity level. A region-based, symmetric geometric transfer matrix framework for PVC was used, processing both PET images and parcel maps to compute a scalar activity estimate for each parcel. The PET images analyzed included CMRglc, CBF, and CMRO_2_.

### 3. Parcellations

Resting state networks (RSNs) are hierarchically organized at multiple levels of granularity (Doucet, Naveau et al. 2011, Gotts, Gilmore et al. 2020). In this study, cortical parcellations were obtained from (Schaefer, Kong et al. 2018) and resampled to the (3mm)^3^ atlas space. For analyses involving generalized additive models (GAMs), we used the 300-parcellation scheme. From these 300 parcels, regions with a low signal-to-noise ratio (SNR), as defined previously on the basis of a mean BOLD signal map (Ojemann, Akbudak et al. 1997), comprising the medial prefrontal cortex and anterior/ventral portions of the temporal lobe, were excluded. These regions were mainly associated with the limbic network as defined in Schaefer’s 17 networks, reducing the number of parcels to 271 for analysis. For comparing spectral slope maps with metabolic measures from the AMBR dataset, which was processed for PVC using the 200-parcellation scheme, we also used the 200-parcellation scheme.

### 4. Power spectral density (PSD) calculation from autocorrelation function

We computed the PSD from the lagged autocovariance function by employing a cosine-based Fourier transform (‘Wiener-Khinchin theorem’) to allow for the exclusion of high-motion frames. First, we computed a scaled temporal lag q associated with each lag m (q = (m-1)*TR/30), where TR represents the sampling interval, and 30 serves as a normalization factor for this analysis. The autocovariance values were adjusted by an exponential decay factor (exp(−0.5q^2^)/n) to account for the decrease in signal similarity with increasing time lags. This exponential decay factor is based on the temporal weighting factor q as well as the number of valid frame pairs n, thereby ensuring the analysis considers only those frames that are relatively *motion-free* (we use the term “motion-free” loosely, as these frames are still subject to head motion, albeit less so than the censored frames). Finally, a cosine transform was applied to the adjusted lagged autocovariance function for each voxel, yielding the PSD for each voxel.

### 5. Spectral slope calculation

BOLD fluctuations exhibit a scale-free activity (He 2011). He and colleagues have previously established that a power-law function, particularly within the <0.1Hz frequency range, more accurately fits the fMRI power spectrum than exponential and log-normal functions (He 2011). Following this work, our prior work used the power-law exponent (Park, Snyder et al. 2023). In the current study, we compared the “fitness” of the spectral slope between fitting the slope in log-log vs. log-normal function by evaluating R-squared values. Log-normal functions consistently yield higher R-squared values compared to those derived from the log-log functions (see **Supplemental Materials**). Accordingly, in the present study, we defined spectral slope as the negative of the first derivative of the slope fitted to the logged power vs. raw frequency values.

### 6. Characterizing age effects

#### 6.1. Generalized Additive Models (GAMs)

Statistics for generalized additive models (GAMs) were performed using RStudio 2023.12.1+402 “Ocean Storm” and R 4.3.3. Drawing from prior work that utilized GAMs to analyze developmental effects (Baum, Flournoy et al. 2022, Sydnor, Larsen et al. 2023), the current study expands on GAMs to characterize the impacts of aging, seen as part of continuing development/lifespan changes. GAMs are advantageous for the flexibility in modeling both linear and non-linear relations between independent and dependent variables. The methodology for GAMs used in this work follows the framework detailed in (Sydnor, Larsen et al. 2023). Briefly, GAMs were fit with parcel-specific spectral slope as the dependent variable, including age as a smooth term and sex and in-scanner head motion as linear covariates. Models for each parcel were evaluated using thin plate regression splines as the smooth term basis set, applying the restricted maximal likelihood approach for smoothing parameters and limiting the maximum basis complexity (k) to 4. The significance of the association between spectral slope and age for each parcel was assessed using an analysis of variance (ANOVA), contrasting the full model (including both age and covariates) with a reduced model (only including covariates). The difference in the explained variance between the full model and the reduced model was denoted as the age-specific R^2^ difference. A significant result indicates a substantial reduction in residual deviation when age is included, as determined using the chi-squared test statistic. The p-values from the ANOVA across all parcel-specific GAMs were adjusted using the FDR correction. The threshold for statistical significance was set at P_FDR_ <0.05.

#### 6.2. Associations with cerebral metabolic rate of glucose (CMRGlc)

We evaluated whether the extent to which spectral slope changed with age correlated spatially with “normative” CMRGlc topography. The “normative” CMRGlc topography was defined by the average baseline CMRGlc of young adults, corrected for PVC across Schaefer’s 200 parcels in the AMBR dataset. The correlation coefficient was computed between the age-specific R^2^ values (comparing the full GAM model against the reduced GAM model) and the “normative” CMRglc map. The association between these two spatial maps was quantified with Spearman correlation, a non-parametric, rank-based method that does not depend on the assumption of normality.

To test the significance of Spearman correlation, we adapted spin permutation tests similar to those described by (Gordon, Laumann et al. 2016, <u>Alexander-Bloch, Shou et al.</u>). Since age-specific R^2^ difference and CMRGlc values were both computed in volumetric space using the Schaefer 200 parcellation scheme, we first resampled the centroid coordinates of these parcellations in surface space for random rotations. Each random rotation, repeated 10,000 times using the quaternion approach (Kuipers 1999), generated distributions of randomly rotated age-specific R2 difference values. The detailed approach is included in the **Supplemental Materials**. In each rotation, parcels rotated into the medial wall were excluded when computing Spearman correlation between the rotated values and parcel-specific CMRGlc values. The p-value from the spin permutation tests, p_spin_, was defined as the number of permuted correlations that were greater than the original Spearman correlations.

#### 6.3. Fuzzy c-means clustering

The fuzzy-c-means clustering analysis was conducted in MATLAB 2023b. The *predict* function in R used for the full GAM model outputs the smooth function for age, which function represents the lifespan trajectory of each parcel’s spectral slope. Observing distinct patterns in each parcel’s lifespan trajectory showing varying levels of linearity, we applied fuzzy c-means clustering to assign a weighted membership between zero and one to each parcel’s trajectory. First, to determine the optimal cluster number, *fcm* was evaluated by incrementally increasing the cluster count (N = 2 - 15). Options used for *fcm* were as follows: the exponent of the fuzzy partition matrix was set to 5 to allow a greater degree of overlap; the maximum iteration count was set to 10,000; the default value for the minimum objective function improvement between two consecutive iterations was used (1e-5); Euclidean distance was used as a distance metric. Subsequently, for each cluster number, a silhouette score was computed by first converting the weighted membership into a binary format using a winner-take-all analysis. Two clusters resulted in the highest silhouette score (see **Supplemental Materials**). Thus, N = 2 was used to cluster the trajectories and their corresponding cortical regions. Subsequently, we evaluated whether clustering based on aging trajectories aligned with well-known resting state networks (RSNs) (Seitzman, Gratton et al. 2020, Dworetsky, Seitzman et al. 2021). Specifically, we leveraged voxelwise RSN membership probability maps (Luckett, Lee et al. 2022) and computed the normalized sum of dot products between RSN probability maps and clustering membership values assigned for all parcels in each cluster.

### 7. Spectral Slope Youthful Index analysis

#### 7.1. Spectral Slope Youthful Index

Goyal et al. introduced the “youthful pattern” analysis, where individual brain metabolism (fludeoxyglucose and oxygen metabolism) and cerebral blood flow (CBF) maps were compared to the corresponding group maps from a young adult cohort using Spearman rank correlation (Goyal, Blazey et al. 2023). Extending this approach, we defined the spectral slope (SS) Youthful Index by evaluating the Spearman correlation between individual SS maps and the averaged SS maps from younger individuals (<45 yrs) in the CamCAN dataset (dataset described in **Methods 1.1**). For outlier analysis, we identified subjects whose SS youthful indices were greater than 3 scaled median absolute deviations from the median across all individuals in the CamCAN dataset. These outliers, represented as red symbols and outlined with a red dashed box in **Fig. 4A**, are also highlighted in **Fig. 4B-D**.

#### 7.2. Spectral Slope topography vs. metabolic and hemodynamic maps

SS maps were additionally compared with averaged metabolic (CMRGlc/CMRO_2_) and hemodynamic (CBF) maps from individuals under 45 years in the AMBR dataset (dataset described in **Methods 1.2**)). The objective of this analysis was twofold: 1) to evaluate changes in SS topography with aging and 2) to assess its similarity and changes in its similarity to “youthful” CMRGlc/CMRO_2_/CBF maps with aging.

#### 7.3. Evaluation of potential confounders in outliers identified by the Spectral Slope Youthful Index

We tested whether outliers identified by the SS Youthful Index were driven by potential confounding factors including grey matter volume, head motion, sex differences, or white matter abnormalities. Considering the greater variability observed among older subjects, we specifically included individuals aged over 45 years. Grey matter volume for each individual was quantified using FSL FAST segmentation, which segments the atlas-registered individual’s T1w image into three tissue types: grey matter, white matter, and CSF. Total individual-specific grey matter volume was defined by summing the number of segmented grey matter voxels in (3mm)^3^ atlas space. Head motion was assessed using the previously described realignment procedure (Park, Snyder et al. 2023). Briefly, this measure averages the changes in head displacement and rotation in quadrature.

White matter abnormalities were assessed using normalized T2-weighted (T2w) image intensities based on an intensity atlas, adopting a technique described in (Brier, Snyder et al. 2021). Brier and colleagues used T1-weighted and FLAIR images. However, as we did not have FLAIR image available, we used T1w and T2w images for image intensity normalization. The reference intensity atlas was computed based on CamCAN participants under 40 years. Each participant’s T2w image was normalized to the reference intensity atlas. To analyze the association between white matter and SSYI on a voxel-by-voxel basis, we implemented an approach similar to voxel-based lesion-symptom mapping (VLSM) (Bates, Wilson et al. 2003). For each voxel, instead of employing conventional group comparisons, we treated SSYI and T2w image intensities as continuous variables for computing Pearson correlations. Considering the exploratory nature of this analysis, we set the significance threshold at α = 0.05. Voxels showing significant correlations with SSYI were subsequently used as an ROI. The averaged T2w image intensity value within this ROI was used as a representative measure of white matter abnormalities.

To quantitatively determine if any of these variables explain the difference between individuals in the outlier vs non-outlier groups, we fitted a linear regression model with the binarized SS Youthful Index as the response variable. Age, grey matter volume (GMV), head motion (HM), and frontal lobe white matter T2w image intensities (FLWM T2w) were used as continuous predictor variables, and sex as a categorical predictor variable:

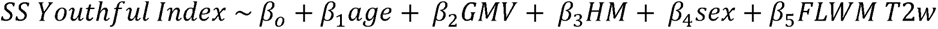

## Acknowledgments and funding sources

Data collection and sharing for this work was provided by the NIH/NIA grants R01 AG053503 (PI: AGV), R01 AG057536 (PIs: MSG, AGV), and RF1 AG073210 (PIs: MSG, AGV), and Cambridge Centre for Ageing and Neuroscience (CamCAN). Funding for CamCAN was provided by the UK Biotechnology and Biological Sciences Research Council under grant number BB/H008217/1 and was additionally supported by the UK Medical Research Council and the University of Cambridge, UK. Parts of this work were funded by the Intellectual and Developmental Disabilities Research Center at Washington University award Number P50 HD103525.

Cognitive, Computational, and Systems Neuroscience (CCSN) Fellowship, McDonnell Center for Systems Neuroscience, Washington University School of Medicine (KYP)

NIH R01 CA203861 (KYP, AZS, JJL, JSS, ECL)

NIH P41 EB018783 (ECL)

NIH U24 NS109103 (ECL)

NIH R01EB026439 (ECL)

NIH/NIA R01 AG053503 (AGV)

NIH/NIA R01 AG057536 (MSG, AGV)

NIH/NIA RF1 AG073210 (MSG, AGV)

## Data Availability

The Cam-CAN dataset is publicly available at https://cam-can.mrc-cbu.cam.ac.uk/dataset/. AMBR dataset: Data availability is based on prior subject consents and the 2018 Common Rule. Coded and processed regional data, suitable for additional data and statistical analyses, can be obtained from the study authors upon reasonable request by a qualified researcher under a data use agreement. For access to raw imaging data, requests should be directed to the VG Lab (https://www.mir.wustl.edu/research/research-centers/neuroimaging-labs-research-center-nil-rc/labs/vlassenko-goyal-lab/), where these data were originally collected. The primary atlas registration pipeline used in this work is available at https://github.com/robbisg/4dfp_tools. ANTs registration can be found at http://stnava.github.io/ANTs/. Any other codes used in this study will be available upon request to KYP.

## Supplementary Materials

### Analyzing BOLD fluctuations in the temporal frequency domain

Prior analyses of BOLD signal fluctuations have largely focused on the amplitude of low-frequency fluctuations (ALFF) (<u>Zang, He et al. 2007</u>) and fractional ALFF (fALFF) (<u>Zou, Zhu et al. 2008</u>). Comparatively, fewer studies have focused on the spectral slope (He 2011, Baria, Mansour et al. 2013). ALFF quantitates the average power within a selected infra-slow frequency range (typically, <0.1 Hz). In contrast, fractional ALFF (fALFF) represents the relative prevalence of slow vs. fast activity. Accordingly, fALFF quantifies the proportion of infra-slow frequency activity relative to the full frequency spectrum (footnote: the range of measurable frequencies depends on the volume TR and fMRI run duration). Spectral slope characterizes the power spectrum in terms of a single number. This is possible because BOLD fluctuations are approximately scale-free, i.e., exhibit a 1/f-like power spectrum (Baria, Mansour et al. 2013, Tagliazucchi, von Wegner et al. 2013). Thus, the SS measure captures information similar to fALFF, although at a finer spectral resolution and utilizing more of the frequency spectrum.

### Log-log vs. log-linear spectral slope fitting

Prior rs-fMRI work modeled BOLD signal spectral slope either as a log-log or log-linear relation (He 2011, Baria, Mansour et al. 2013, Park, Snyder et al. 2023). In the present study, we first compared the spectral slope fit using both approaches: a log-log analysis where both power and frequency were logarithmically transformed; and a log-linear analysis where only power was logarithmically transformed. For each method, a linear model described the relations between frequency and power. Specifically, the spectral slope was computed as the negative of the first derivative of the fitted slope.

To measure the model accuracy, we computed the coefficient of determination (R^2^), which computes the proportion of variance in the dependent variable predictable from the independent variable. This was calculated by first summing the squared deviations between the fitted slopes and the actual spectra to obtain the residual sum of squares, which indicates the variance unexplained by the model. The total sum of squares was computed by summing the squared deviations of the actual power spectra values from their mean, reflecting the total variance observed. R^2^ was derived by subtracting the ratio of residual sum of squares to the total sum of squares from one, with values closer to 1 indicating a more accurate model.

R^2^ values were consistently higher with the log-linear models, suggesting a better fit compared to the log-log approach (**Fig. S1**). Based on these findings, we used the log-linear slope fitting for our spectral slope metric.

**Figure S1.**
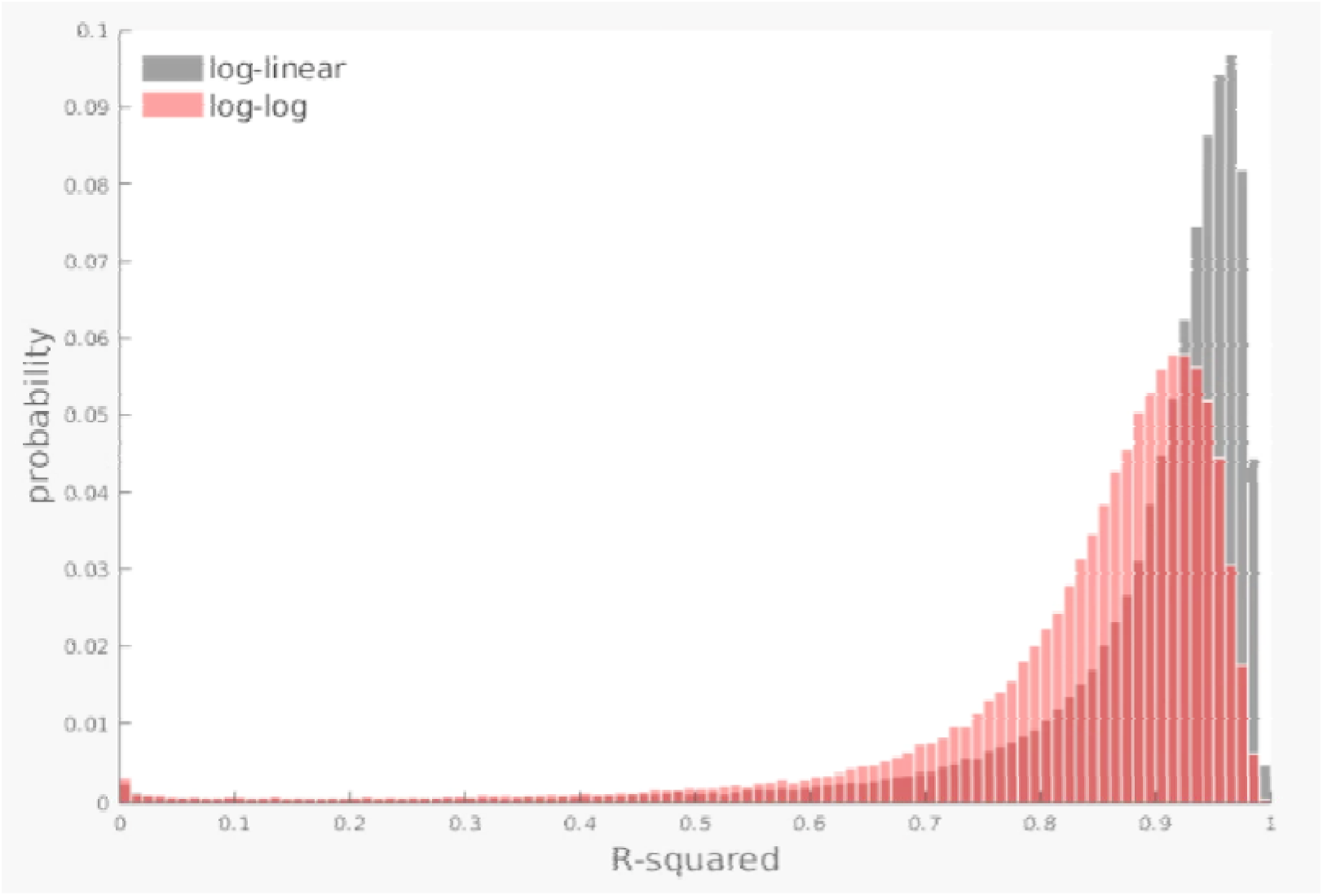
Distribution of R^2^ values of spectral slope fits in log-linear and log-log domains. The log-linear domain (gray) consistently shows higher R^2^ values compared to the log-log domain (red).

### Spin permutation test: rotation quaternions

To test the significance of any model, it is useful to generate a null hypothesis. Here, the null hypothesis is no difference in the spatial similarity between the age effect in the spatial topography of spectral slope or CMRGlc. It is necessary to include spatial autocorrelation in the construction of the null model. Prior methodology to construct an appropriate null model, i.e., the spin test (Gordon, Laumann et al. 2016, Alexander-Bloch, Shou et al.), generated surrogate data by randomly rotating brain surface distributions represented on the inflated sphere. In detail, the distribution of rotations achieved by this prior version of the spin test was not uniform. The present methodology achieves a uniform distribution of rotations by sampling random quaternions.

A quaternion is defined as four components: one real and three imaginary parts. These components are computed using the following formula:

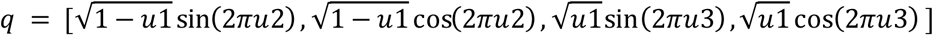

where u1, u2, and u3 are uniformly distributed random numbers between 0 and 1. Note that 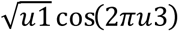 is the real part of the quaternion and the remaining components are the vector part of the quaternion. This formulation ensures a uniform distribution in the rotation group *SO*(3).

To apply the rotation, each quaternion is converted into a corresponding 3×3 rotation matrix, R:

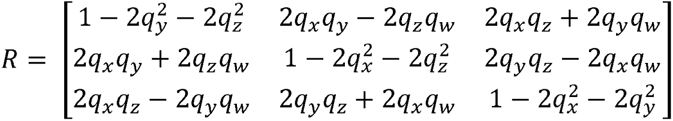

The variables *q_w_*, *q_x_*, *q_y_*, and *q_z_* represent the real and imaginary parts of quaternion, respectively. This matrix applies the quaternion’s defined spatial orientation to rotate each vertex. (See (Kuipers 1999) Chapter 5, Section 5.14 “Quaternions to Matrices” for more details.)

The remaining operations are performed as described in (Gordon, Laumann et al. 2016, Alexander-Bloch, Shou et al.). In brief, both the centroid vertices and the corresponding age-specific R^2^ difference values are reflected across the Y-Z plane. The quaternion used for the left hemisphere is also applied to the right hemisphere for each random sample rotation. Centroid vertices rotated into the medial wall are excluded when computing the Spearman’s correlation between the rotated values and CMRGlc values.

### Fuzzy Silhouette score analysis

Trajectories in age-related spectral slope flattening were not uniform (main text **Fig. 2**). Accordingly, we performed a silhouette score analysis (Rousseeuw 1987) using the fuzzy c-means clustering algorithm to determine the optimal number of clusters. Fuzzy c-means clustering was applied to z-scored GAM smooth functions that chart the trajectory of SS changes with age, across a range of 2 to 15 clusters.

Silhouette score analysis requires hard cluster assignments. Accordingly, we first converted fuzzy membership scores to hard assignments by assigning the parcel to the cluster with the highest membership value. Subsequently, we computed the silhouette scores for each parcel using the Euclidean distance metric. The silhouette value of each parcel quantifies its similarity to other parcels within its own cluster compared to parcels in the nearest cluster. The average silhouette score for each cluster number was computed by averaging the silhouette scores across all parcels (**Fig. S2**). The final result showed that the optimal number of clusters was 2.

**Figure S2.**
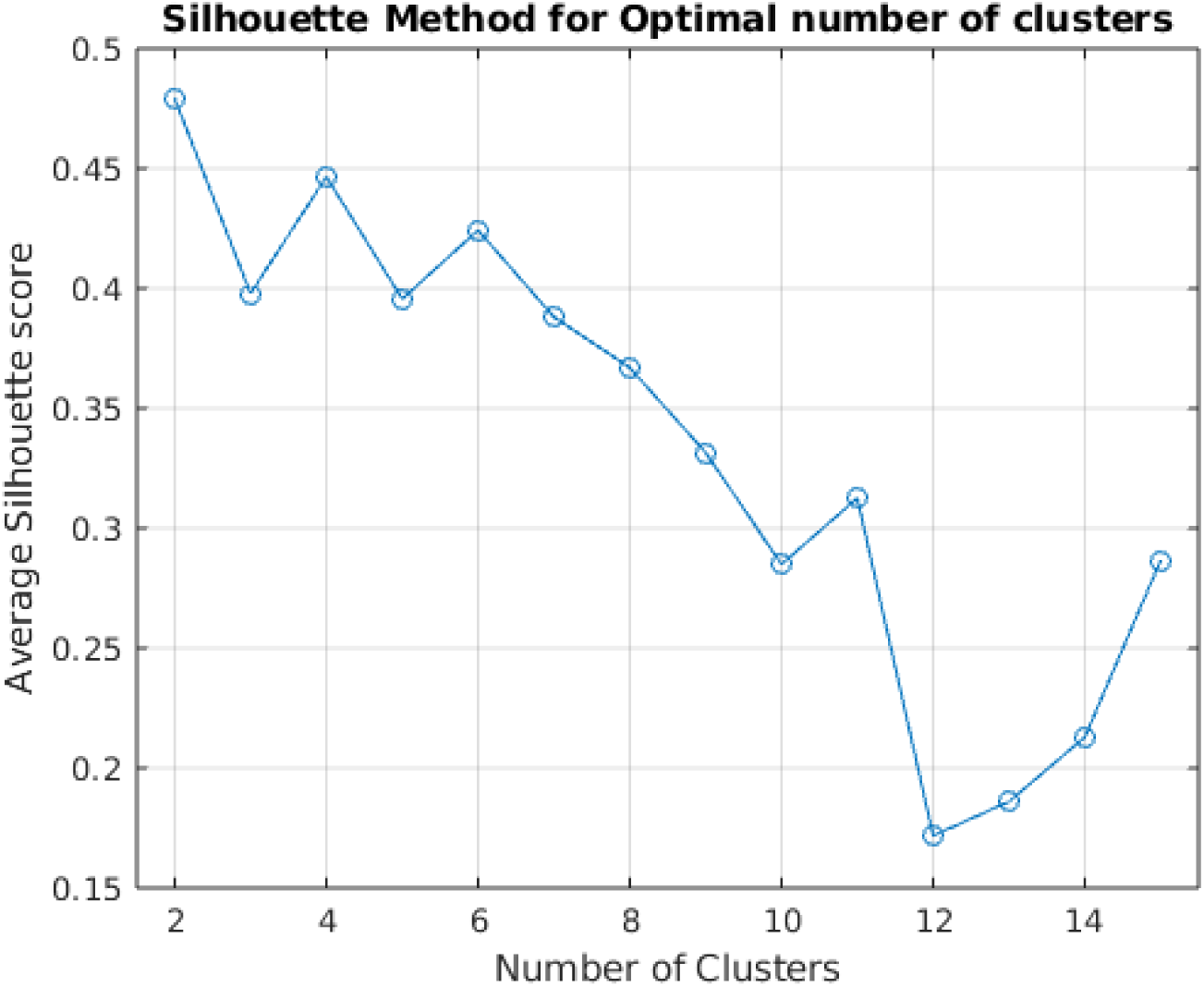
Average silhouette scores for each cluster number, ranging from 2 to 15. Higher silhouette scores indicate better cluster cohesion and separation.

### Linear regression model result

**Supplementary Table 1.**
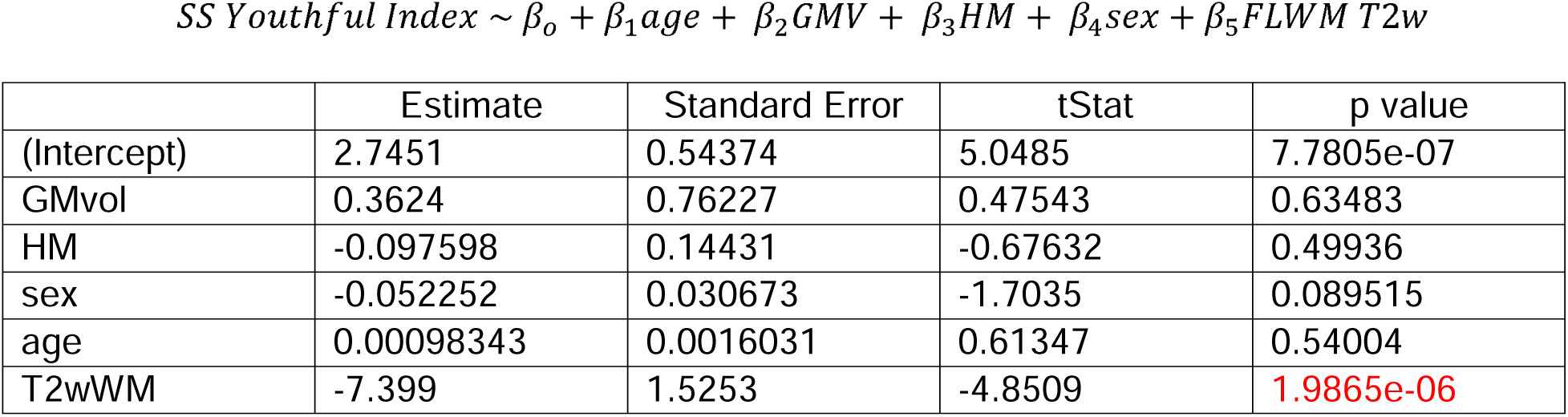
Relation between outlier status and frontal lobe white matter intensity (p < 2e-06). Linear regression model (*fitlm* function in MATLAB 2023b) results including grey matter volume (GMvol), head motion (HM), sex, age, and T2-weighted frontal lobe white matter intensity as independent variables, and outlier status as the dependent categorical variable. See main text Figure 5. Note that only subjects older than 45 years is included in this analysis.

**Supplementary Table 2.**
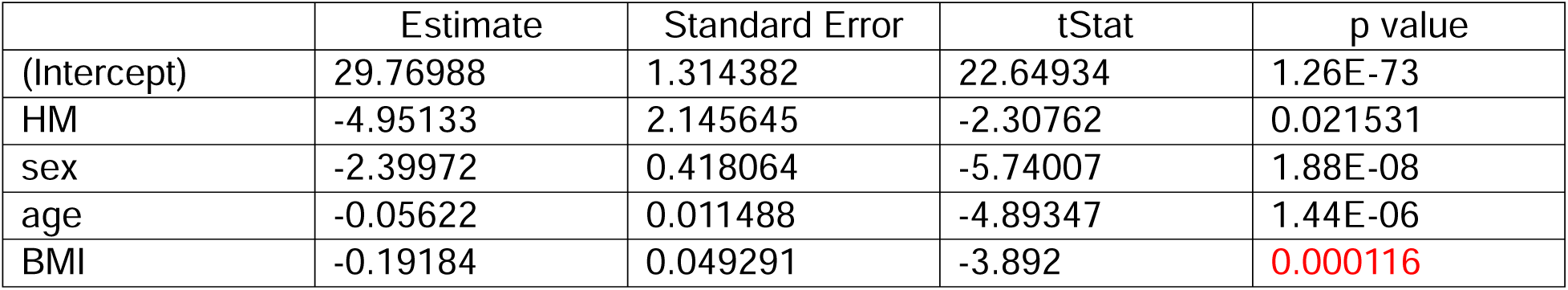
Relation between whole-brain spectral slope average and BMI (p <1.2e-04). Linear regression model (*fitlm* function in MATLAB 2023b) results including head motion (HM), sex, age, and BMI as independent variables, and whole-brain spectral slope average as a dependent variable. See Figure S3 (left panel).

**Supplementary Table 3.**
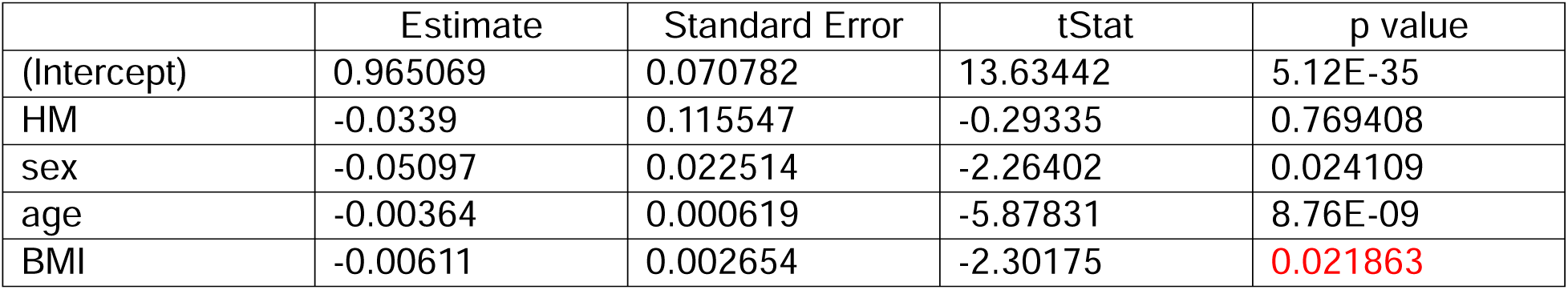
Relation between spectral slope youthful index (SSYI) and BMI (p < 0.03). Linear regression model (*fitlm* function in MATLAB 2023b) results including head motion (HM), sex, age, and BMI as independent variables, and SSYI as a dependent variable. See Figure S3 (right panel).

**Figure S3.**
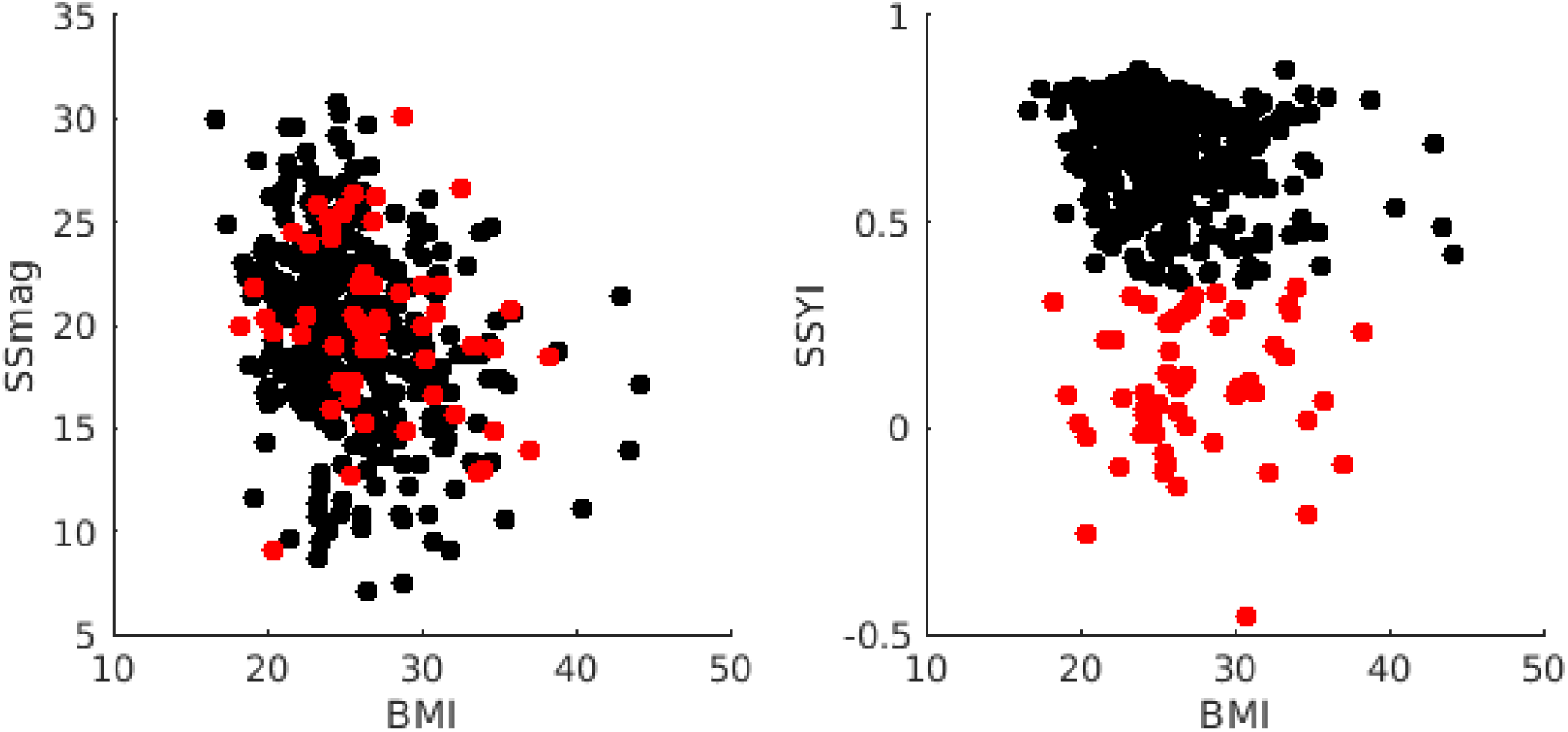
Relation between BMI and whole-brain spectral slope average (SSmag) and SSYI. Outliers and non-outliers, as shown in Figures 4 and 5 of the main text, are represented by red and black dots, respectively. Spectral slope measures correlate BMI. This relation is statistically significant after accounting for covariates including age, sex, and head motion (Tables S2 and 3).

### T2w frontal lobe white matter intensities are correlated with body mass index (BMI)

Given that white matter abnormalities are associated with hypertension (Dufouil, de Kersaint-Gilly et al. 2001), we asked whether cardiovascular measures of subjects were related to T2w frontal lobe white matter intensities. We utilized the physiological data provided by CamCAN, which includes systolic blood pressure, diastolic blood pressure, weight, and height. Pulse pressure is calculated as the difference between systolic and diastolic blood pressure. We computed body mass index (BMI) as weight (kg) divided by the square of height (m^2^). BMI is a crude measure of body fat percentage and despite questions regarding the accuracy of BMI as an indicator of cardiovascular health, it is often considered a risk factor for cardiovascular disease (Brown, Higgins et al. 2000). Our findings suggest a significant positive correlation between T2w FLWM intensities and BMI (r = 0.354, p < 1e-06; **Fig. S4**), but no such correlation with pulse pressure.

**Figure S4.**
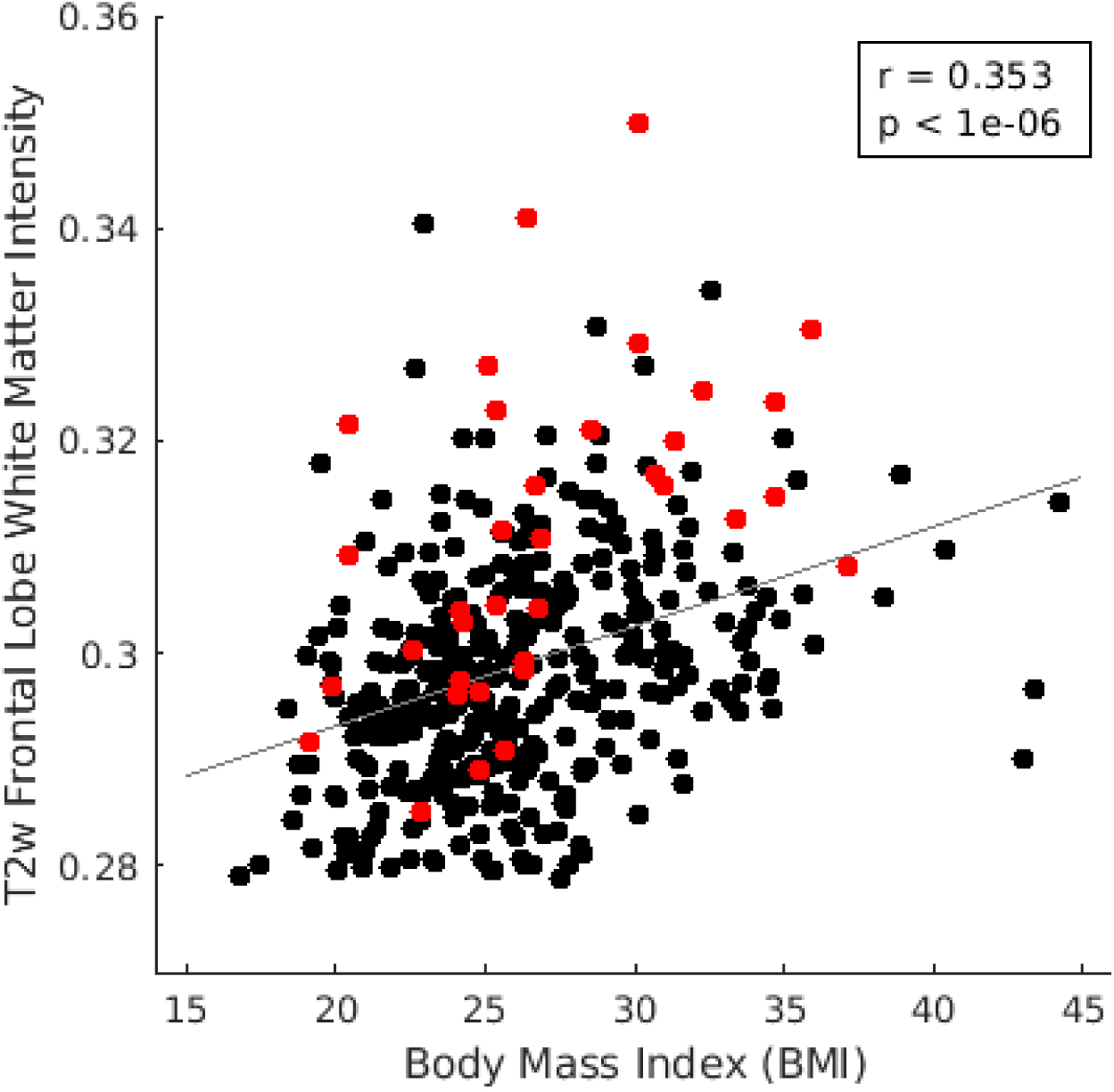
Scatterplot demonstrating the relationship between body mass index (BMI) and T2-weighted (T2w) frontal lobe white matter (FLWM) intensity. Outliers and non-outliers, as shown in Figures 4 and 5 of the main text, are represented by red and black dots, respectively. Note the positive correlation between BMI and T2w FLWM intensity.

