## Supplemental Materials for "Aging and the Spectral Properties of Brain Hemodynamics"

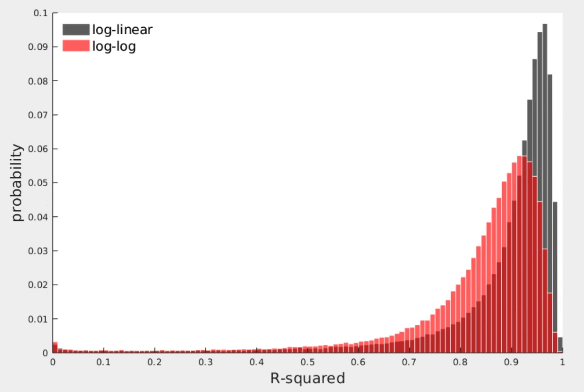

**Figure S1**. Distribution of R^2^ values of spectral slope fits in log-linear and log-log domains. The log-linear domain (gray) consistently shows higher R^2^ values compared to the log-log domain (red).

**Spin permutation test: rotation quaternions**

To test the significance of any model, it is useful to generate a null hypothesis. Here, the null hypothesis is no difference in the spatial similarity between the age effect in the spatial topography of spectral slope or CMRGlc. It is necessary to include spatial autocorrelation in the construction of the null model. Prior methodology to construct an appropriate null model, i.e., the spin test (Gordon, Laumann et al. 2016, Alexander-Bloch, Shou et al.), generated surrogate data by randomly rotating brain surface distributions represented on the inflated sphere. In detail, the distribution of rotations achieved by this prior version of the spin test was not uniform. The present methodology achieves a uniform distribution of rotations by sampling random quaternions.

A quaternion is defined as four components: one real and three imaginary parts. These components are computed using the following formula:

$$q = [\sqrt{1-u1}\sin\left( 2\pi u2 \right),\sqrt{1-u1}\cos\left( 2\pi u2 \right),\sqrt{u1}\sin\left( 2\pi u3 \right),\sqrt{u1}\cos\left( 2\pi u3 \right) ]$$

where u1, u2, and u3 are uniformly distributed random numbers between 0 and 1. Note that $\sqrt{u1}\cos\left( 2\pi u3 \right)$ is the real part of the quaternion and the remaining components are the vector part of the quaternion. This formulation ensures a uniform distribution in the rotation group $SO(3)$.

To apply the rotation, each quaternion is converted into a corresponding 3x3 rotation matrix, R:

$$R= \left[ \begin{matrix} 1-{2q}_{y}^{2}-{2q}_{z}^{2} & {2q}_{x}q_{y}-{2q}_{z}q_{w} & {2q}_{x}q_{z}+{2q}_{y}q_{w} \\ {2q}_{x}q_{y}+{2q}_{z}q_{w} & 1-{2q}_{x}^{2}-{2q}_{z}^{2} & {2q}_{y}q_{z}-{2q}_{x}q_{w} \\ {2q}_{x}q_{z}-{2q}_{y}q_{w} & {2q}_{y}q_{z}+{2q}_{x}q_{w} & 1-{2q}_{x}^{2}-{2q}_{y}^{2} \end{matrix} \right]$$

The variables $q_{w}$, $q_{x}$, $q_{y}$, and $q_{z}$ represent the real and imaginary parts of quaternion $q$, respectively. This matrix applies the quaternion’s defined spatial orientation to rotate each vertex. (See (Kuipers 1999) Chapter 5, Section 5.14 “Quaternions to Matrices” for more details.)

The remaining operations are performed as described in (Gordon, Laumann et al. 2016, Alexander-Bloch, Shou et al.). In brief, both the centroid vertices and the corresponding age-specific R^2^ difference values are reflected across the Y-Z plane. The quaternion used for the left hemisphere is also applied to the right hemisphere for each random sample rotation. Centroid vertices rotated into the medial wall are excluded when computing the Spearman’s correlation between the rotated values and CMRGlc values.

**Fuzzy Silhouette score analysis**

Trajectories in age-related spectral slope flattening were not uniform (main text **Fig. 2**). Accordingly, we performed a silhouette score analysis (Rousseeuw 1987) using the fuzzy c-means clustering algorithm to determine the optimal number of clusters. Fuzzy c-means clustering was applied to z-scored GAM smooth functions that chart the trajectory of SS changes with age, across a range of 2 to 15 clusters.

Silhouette score analysis requires hard cluster assignments. Accordingly, we first converted fuzzy membership scores to hard assignments by assigning the parcel to the cluster with the highest membership value. Subsequently, we computed the silhouette scores for each parcel using the Euclidean distance metric. The silhouette value of each parcel quantifies its similarity to other parcels within its own cluster compared to parcels in the nearest cluster. The average silhouette score for each cluster number was computed by averaging the silhouette scores across all parcels (**Fig. S2**). The final result showed that the optimal number of clusters was 2.

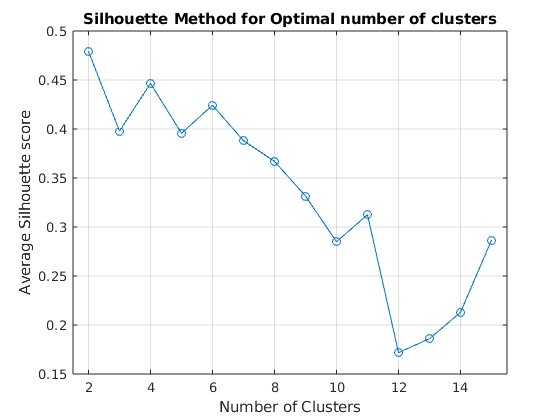

**Figure S2**. Average silhouette scores for each cluster number, ranging from 2 to 15. Higher silhouette scores indicate better cluster cohesion and separation.

**Linear regression model result**

**Supplementary Table 1.** Relation between outlier status and frontal lobe white matter intensity (p < 2e-06). Linear regression model (*fitlm* function in MATLAB 2023b) results including grey matter volume (GMvol), head motion (HM), sex, age, and T2-weighted frontal lobe white matter intensity as independent variables, and outlier status as the dependent categorical variable. See main text Figure 5. Note that only subjects older than 45 years is included in this analysis.

$$SS Youthful Index \sim\beta_{o}+\beta_{1}age+ \beta_{2}GMV+ \beta_{3}HM+ \beta_{4}sex+\beta_{5}FLWM T2w$$

|  | Estimate | Standard Error | tStat | p value |
| --- | --- | --- | --- | --- |
| (Intercept) | 2.7451 | 0.54374 | 5.0485 | 7.7805e-07 |
| GMvol | 0.3624 | 0.76227 | 0.47543 | 0.63483 |
| HM | -0.097598 | 0.14431 | -0.67632 | 0.49936 |
| sex | -0.052252 | 0.030673 | -1.7035 | 0.089515 |
| age | 0.00098343 | 0.0016031 | 0.61347 | 0.54004 |
| T2wWM | -7.399 | 1.5253 | -4.8509 | 1.9865e-06 |

**Supplementary Table 2.** Relation between whole-brain spectral slope average and BMI (p <1.2e-04). Linear regression model (*fitlm* function in MATLAB 2023b) results including head motion (HM), sex, age, and BMI as independent variables, and whole-brain spectral slope average as a dependent variable. See Figure S3 (left panel).

|  | Estimate | Standard Error | tStat | p value |
| --- | --- | --- | --- | --- |
| (Intercept) | 29.76988 | 1.314382 | 22.64934 | 1.26E-73 |
| HM | -4.95133 | 2.145645 | -2.30762 | 0.021531 |
| sex | -2.39972 | 0.418064 | -5.74007 | 1.88E-08 |
| age | -0.05622 | 0.011488 | -4.89347 | 1.44E-06 |
| BMI | -0.19184 | 0.049291 | -3.892 | 0.000116 |

**Supplementary Table 3.** Relation between spectral slope youthful index (SSYI) and BMI (p < 0.03). Linear regression model (*fitlm* function in MATLAB 2023b) results including head motion (HM), sex, age, and BMI as independent variables, and SSYI as a dependent variable. See Figure S3 (right panel).

|  | Estimate | Standard Error | tStat | p value |
| --- | --- | --- | --- | --- |
| (Intercept) | 0.965069 | 0.070782 | 13.63442 | 5.12E-35 |
| HM | -0.0339 | 0.115547 | -0.29335 | 0.769408 |
| sex | -0.05097 | 0.022514 | -2.26402 | 0.024109 |
| age | -0.00364 | 0.000619 | -5.87831 | 8.76E-09 |
| BMI | -0.00611 | 0.002654 | -2.30175 | 0.021863 |

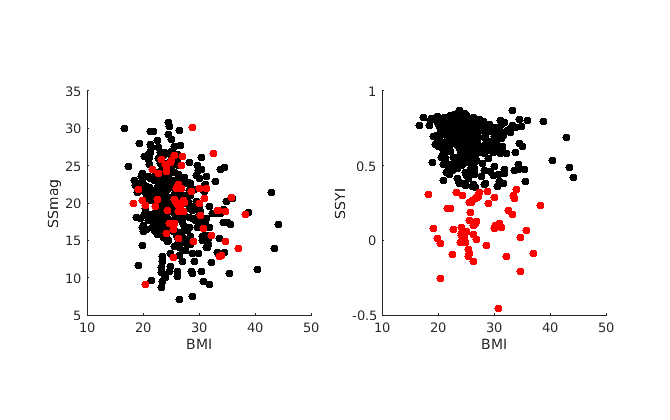

**Figure S3.** Relation between BMI and whole-brain spectral slope average (SSmag) and SSYI. Outliers and non-outliers, as shown in Figures 4 and 5 of the main text, are represented by red and black dots, respectively. Spectral slope measures correlate BMI. This relation is statistically significant after accounting for covariates including age, sex, and head motion (Tables S2 and 3).

**T2w frontal lobe white matter intensities are correlated with body mass index (BMI)**

Given that white matter abnormalities are associated with hypertension (Dufouil, de Kersaint-Gilly et al. 2001), we asked whether cardiovascular measures of subjects were related to T2w frontal lobe white matter intensities. We utilized the physiological data provided by CamCAN, which includes systolic blood pressure, diastolic blood pressure, weight, and height. Pulse pressure is calculated as the difference between systolic and diastolic blood pressure. We computed body mass index (BMI) as weight (kg) divided by the square of height (m^2^). BMI is a crude measure of body fat percentage and despite questions regarding the accuracy of BMI as an indicator of cardiovascular health, it is often considered a risk factor for cardiovascular disease (Brown, Higgins et al. 2000). Our findings suggest a significant positive correlation between T2w FLWM intensities and BMI (r = 0.354, p < 1e-06; **Fig. S4**), but no such correlation with pulse pressure.

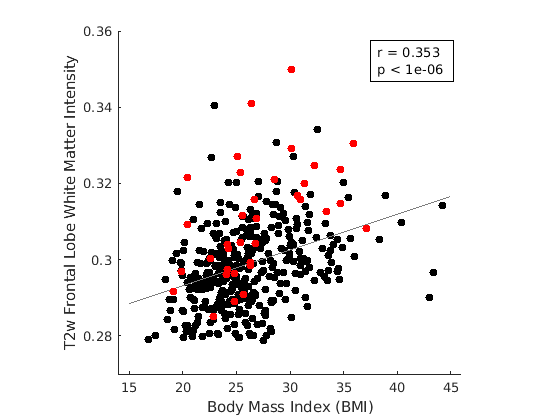

**Figure S4**. Scatterplot demonstrating the relationship between body mass index (BMI) and T2-weighted (T2w) frontal lobe white matter (FLWM) intensity. Outliers and non-outliers, as shown in Figures 4 and 5 of the main text, are represented by red and black dots, respectively. Note the positive correlation between BMI and T2w FLWM intensity.

Alexander-Bloch, A. F., H. Shou, S. Liu, T. D. Satterthwaite, D. C. Glahn, R. T. Shinohara, S. N. Vandekar and A. Raznahan (2018). "On testing for spatial correspondence between maps of human brain structure and function." Neuroimage **178**: 540-551.

Baria, A. T., A. Mansour, L. Huang, M. N. Baliki, G. A. Cecchi, M. M. Mesulam and A. V. Apkarian (2013). "Linking human brain local activity fluctuations to structural and functional network architectures." NeuroImage **73**: 144-155.

Brown, C. D., M. Higgins, K. A. Donato, F. C. Rohde, R. Garrison, E. Obarzanek, N. D. Ernst and M. Horan (2000). "Body mass index and the prevalence of hypertension and dyslipidemia." Obes Res **8**(9): 605-619.

Dufouil, C., A. de Kersaint-Gilly, V. Besancon, C. Levy, E. Auffray, L. Brunnereau, A. Alperovitch and C. Tzourio (2001). "Longitudinal study of blood pressure and white matter hyperintensities: the EVA MRI Cohort." Neurology **56**(7): 921-926.

Gordon, E. M., T. O. Laumann, B. Adeyemo, J. F. Huckins, W. M. Kelley and S. E. Petersen (2016). "Generation and Evaluation of a Cortical Area Parcellation from Resting-State Correlations." Cereb Cortex **26**(1): 288-303.

He, B. J. (2011). "Scale-Free Properties of the Functional Magnetic Resonance Imaging Signal during Rest and Task." Journal of Neuroscience **31**(39): 13786-13795.

Kuipers, J. B. (1999). Quaternions and rotation sequences : a primer with applications to orbits, aerospace, and virtual reality. Princeton, N.J., Princeton University Press.

Park, K. Y., A. Z. Snyder, M. Olufawo, G. Trevino, P. H. Luckett, B. Lamichhane, T. Xie, J. J. Lee, J. S. Shimony and E. C. Leuthardt (2023). "Glioblastoma induces whole-brain spectral change in resting state fMRI: Associations with clinical comorbidities and overall survival." NeuroImage: Clinical **39**: 103476.

Rousseeuw, P. J. (1987). "Silhouettes: A graphical aid to the interpretation and validation of cluster analysis." Journal of Computational and Applied Mathematics **20**: 53-65.
